# Auditory Working Memory in Adolescents with Specific Learning Disorders

**DOI:** 10.1101/2025.10.16.682902

**Authors:** Yeganeh Salimi, Marjan Makhsous, Ehsan Rezayat

## Abstract

Specific Learning Disorders (SLD), including dyslexia, dyscalculia, and dysgraphia, are associated with deficits in executive functions such as auditory working memory (AWM). This study investigated AWM performance and metacognitive monitoring in adolescents diagnosed with SLD using an auditory delayed-match-to-sample task. Thirty-six participants (18 with SLD and 18 neurotypical controls) were assessed on accuracy, reaction time, and confidence ratings.

Adolescents with SLD exhibited reduced sensitivity to auditory intensity differences, slower reaction times, and more conservative decision-making strategies. Drift diffusion modeling revealed lower evidence accumulation rates and wider decision boundaries in the SLD group. Moreover, confidence ratings were less influenced by stimulus differences, particularly among participants with dyslexia.

These findings highlight impairments in auditory processing, decision-making, and metacognitive self-monitoring in adolescents with SLD. The results underscore the importance of interventions that address not only linguistic skills but also auditory decision-making and metacognitive strategies to support learning outcomes.

## Introduction

Specific Learning Disorders (SLD), including dyslexia, dyscalculia, and dysgraphia, are neurodevelopmental conditions characterized by persistent difficulties in reading, writing, or arithmetic despite adequate intelligence, motivation, and educational opportunities (American Psychiatric Association, 2013). Beyond domain-specific impairments, accumulating evidence indicates that SLD involves broader executive dysfunctions, particularly in working memory (WM), attentional control, and metacognitive regulation (Daniel et al., 2022; Shyam & Venkatesan, 2021).

Working memory, the system responsible for temporarily storing and manipulating information, is critical for learning across modalities. While deficits in visual and verbal WM are well documented in SLD (Habib et al., 2019), auditory working memory (AWM) has received comparatively less attention despite its pivotal role in phonological decoding, language comprehension, and numerical reasoning. AWM supports the retention and comparison of auditory stimuli over short intervals, processes that are fundamental to everyday learning activities such as following multi-step instructions or performing mental calculations (Kumar et al., 2016). Recent studies also indicate that reading disabilities are associated with instability in auditory discrimination thresholds, especially in tasks involving frequency and intensity judgments (McWeeny et al., 2024; Breadmore et al., 2024).

Neurocognitive findings suggest that AWM deficits in SLD may stem from atypical activation of the temporoparietal junction, inferior frontal gyrus, and auditory cortices, regions essential for phonological encoding and auditory attention (Faedda et al., 2019; Czajeczny et al., 2023). Electrophysiological evidence further points to delayed and attenuated P300 responses during auditory discrimination, indicating inefficiencies in encoding and executive filtering of auditory input (Kashani & Tafti, 2018). Together, these studies suggest that AWM difficulties in SLD reflect a combination of sensory-perceptual and higher-order executive dysfunctions.

Beyond task performance, metacognitive monitoring offers a complementary perspective on auditory memory. Confidence ratings serve as indices of self-monitoring and awareness of performance, yet research shows that such metacognitive insight is often compromised in SLD. For example, students with learning disabilities show weaker calibration between accuracy and confidence compared with typically developing peers, which may undermine self-regulation and persistence in academic contexts (Okur & Aksoy, 2025; Daniel et al., 2022). Nevertheless, the relationship between confidence judgments and auditory memory performance across SLD subtypes remains poorly understood. Prior research has primarily focused on phoneme perception and frequency discrimination in dyslexia, linking auditory resolution deficits to phonological difficulties (Banai & Ahissar, 2006; Snowling, 2020). In contrast, intensity discrimination tasks that parametrically manipulate auditory working memory load remain largely unexplored.

The present study addresses this gap by examining auditory working memory and confidence judgments in adolescents with and without SLD. Using a delayed auditory comparison task, we evaluated accuracy, reaction time, and confidence ratings across three subtypes—dyslexia, dyscalculia, and dysgraphia—in comparison with typically developing peers. By integrating behavioral measures with metacognitive assessments and computational modeling, this study aims to (a) characterize the extent of AWM deficits in SLD, (b) assess subtype-specific differences in metacognitive monitoring, and (c) identify cognitive markers with potential diagnostic and educational relevance. Importantly, by combining psychometric and mechanistic approaches, this work provides novel insight into how perceptual, decisional, and metacognitive processes jointly contribute to the learning challenges observed in SLD.

## Materials and methods

### Participants

Forty-four participants were initially recruited for this study. Data from eight individuals were identified as outliers and excluded from subsequent analyses. Participants included 18 individuals diagnosed with Specific Learning Disorders (SLD) and 18 typically developing controls. Within the SLD cohort, 5 participants were classified with dyslexia, 5 with dyscalculia, and 8 with dysgraphia, based on formal clinical diagnosis and corroborated parent interviews. Participants were aged between 10 and 16 years (Mean age = 12.57). The study protocol was reviewed and approved by the Research Ethics Committee of the Faculty of Psychology and Education at the University of Tehran (Approval ID: PSYE-202508-1202), ensuring compliance with ethical standards. Written informed consent was obtained from the parents of all participants, who were informed that their child could withdraw from the study at any time without consequence

### Study Design

This investigation employed a cross-sectional comparative design, involving two independent groups: healthy participants and SLD patients. Demographic information was collected from all participants prior to the experimental sessions. The primary experimental task was a parametric auditory working memory assessment (Akrami et al., 2018).

### Parametric auditory working memory test

Auditory working memory was assessed using pairs of pink noise stimuli presented at varying intensities (60, 62, 64, 66, 68, 70, or 72 dB) through noise-cancelling headphones in a randomized order. Participants completed six practice trials after reviewing instructions to familiarize themselves with the procedure. During each trial, the first stimulus was presented alongside a green square on the left side of the screen. Following a delay of either 2 or 6 seconds, indicated by the word “WAIT!” on the screen, the second stimulus appeared with a red square on the right side. Participants indicated which sound was louder by clicking on the corresponding square using a mouse. Immediately after each response, but before receiving feedback, participants rated their confidence in their choice on a scale from 0 (not confident at all) to 100 (completely confident). Each participant completed 150 trials in total. The task was implemented in MATLAB (The MathWorks Inc., Massachusetts, USA) using Psychtoolbox (Figure 1).

**Figure 1:**
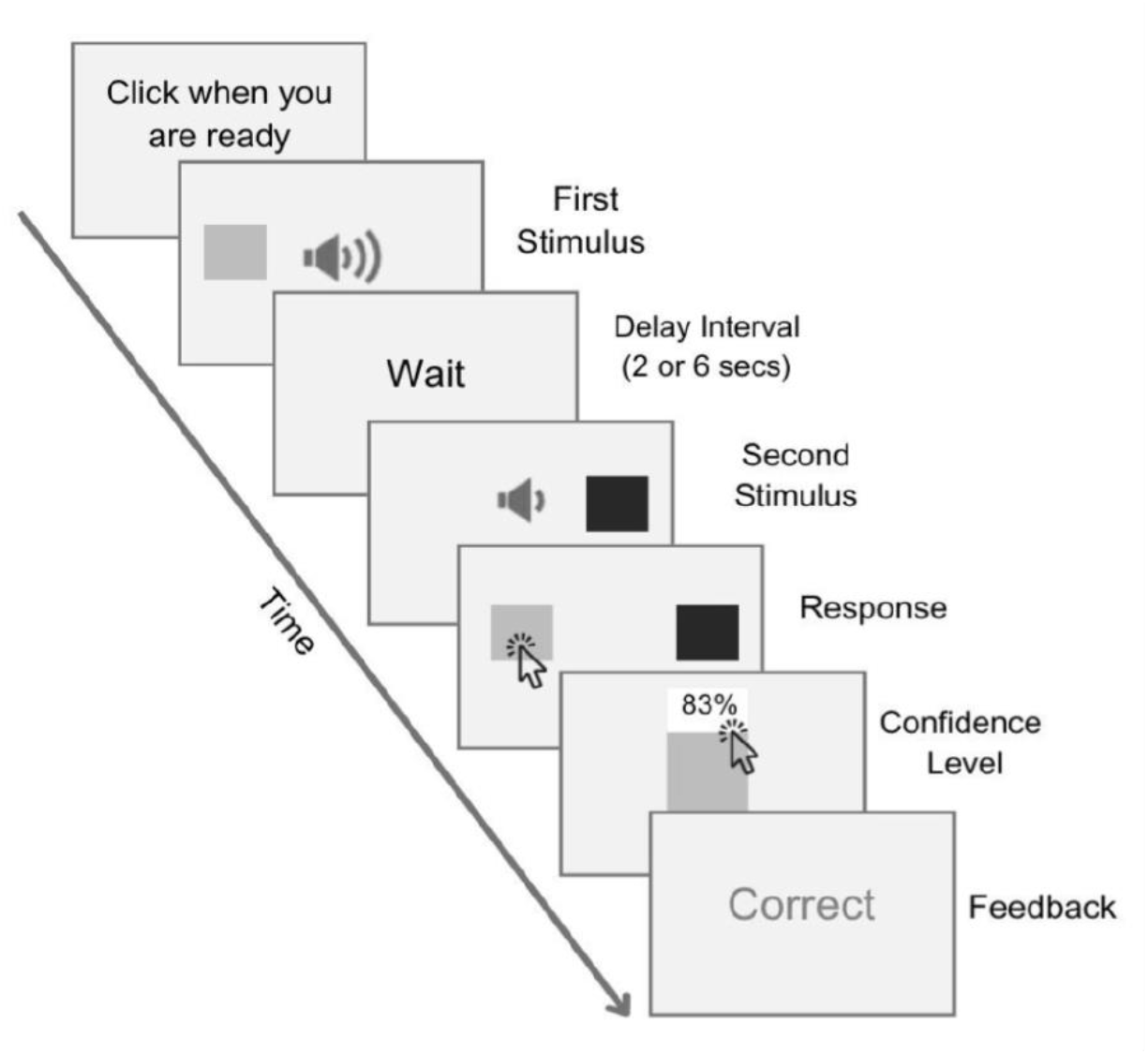
Parametric Auditory Working Memory task. The figure shows the auditory delayed comparison task. Participants compared the loudness of two pink noise sounds, presented with a delay and marked by colored squares. They indicated their choice and confidence, with feedback provided after each of the trials.

### Statistical Analysis

All statistical analyses were conducted in MATLAB, with a significance threshold set at p < 0.05. Individual performance on the auditory working memory task was modeled using binomial generalized linear models (GLMs) with a logit link to estimate psychometric parameters (intercept, slope, threshold, and sensitivity) for each participant across conditions (Figure 2).

**Figure 2:**
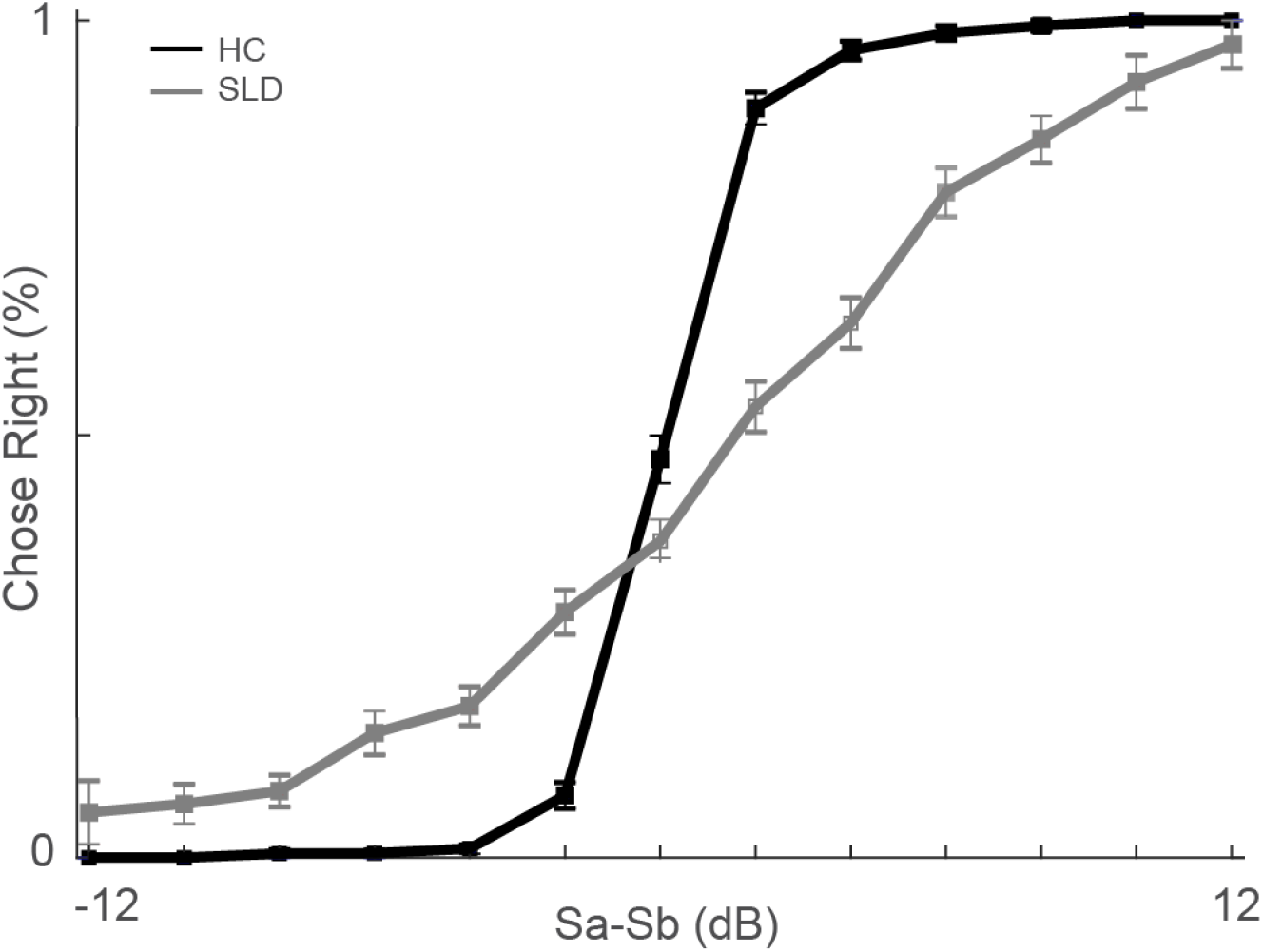
Psychometric functions of auditory discrimination. The figure shows psychometric curves for both groups, depicting the probability of choosing the second stimulus (rightward) based on the dB difference between the first and second stimuli. The y-axis represents the percentage of rightward choices, and the x-axis indicates the stimuli difference.

**Figure 3:**
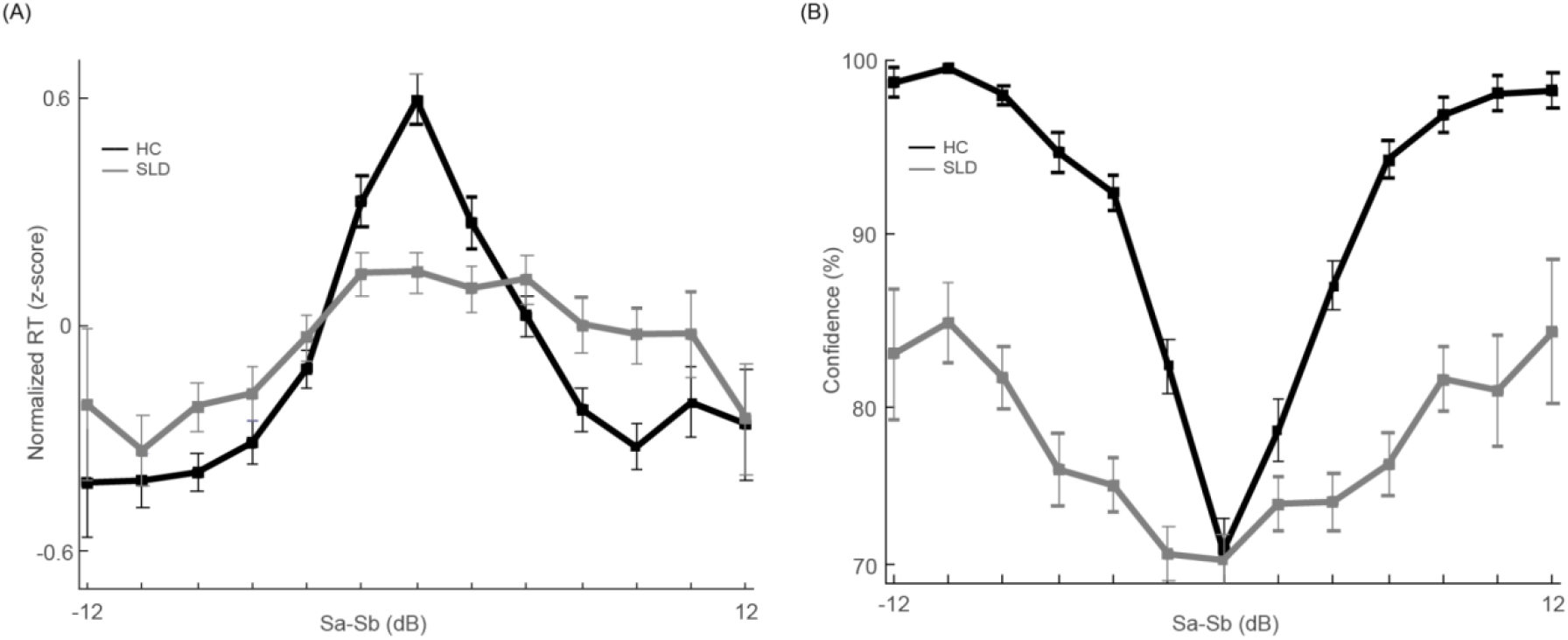
Reaction times and confidence ratings as a function of stimulus intensity differences. Normalized response times (A) and confidence ratings (B) are plotted across levels of stimulus intensity difference (Sa–Sb, in dB) for both groups exhibiting the expected U-shaped pattern in reaction times and monotonic increases in confidence with greater intensity differences. The y-axis represents the normalized response times in A and the percentage of confidence level in B, and the x-axis indicates the stimuli difference in both parts.

**Figure 4:**
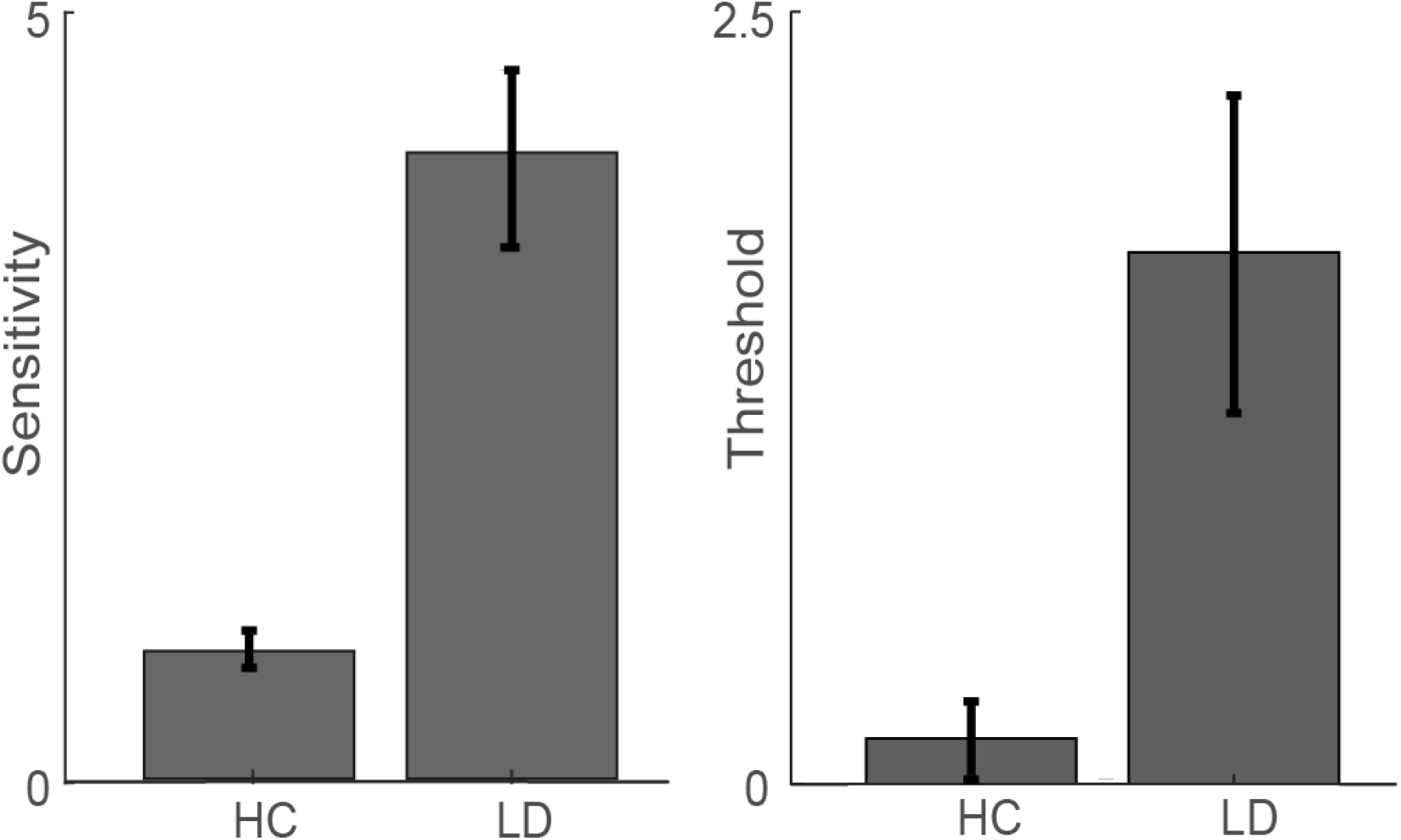
Group differences in psychometric sensitivity and discrimination thresholds. Bar plots show mean sensitivity (left) and discrimination thresholds (right) for healthy controls (HC) and individuals with specific learning disorders (SLD).

Group-level analyses of threshold, sensitivity, log-transformed response times, and confidence ratings were performed using linear mixed-effects models (LMEs). Each model included fixed effects for group, sound intensity differences, SLD type, and delay duration. To account for individual variability, models included random intercepts for participants and random slopes for the difference between sound intensities (1 + ΔSound | participant). Trial-level accuracy (binary outcome) was analyzed using generalized linear mixed-effects models with the same fixed and random effects structure.

In addition, to capture latent decision-making dynamics, we implemented a Drift Diffusion Model (DDM) using the brms package in R with cmdstanr as the backend. The model simultaneously estimated drift rate, boundary separation, starting point bias, and non-decision time. Group-level effects were specified on drift rate, boundary separation, and bias parameters, while non-decision time was modeled with an intercept only. Weakly informative priors, providing reasonable bounds without overly constraining parameter estimates, were applied to all parameters. Models were fit using Hamiltonian Monte Carlo with two chains and 3000 iterations per chain (1500 warm-up). Model convergence was evaluated via R-hat statistics and effective sample sizes, and posterior summaries were used to compare group-level differences in decision parameters.

## Results

A total of 36 participants were included in the analyses, assigned to either the healthy control group (n = 18) or the SLD group (n = 18). Table 1 summarizes the demographic characteristics of the participants.

**Table 1.**
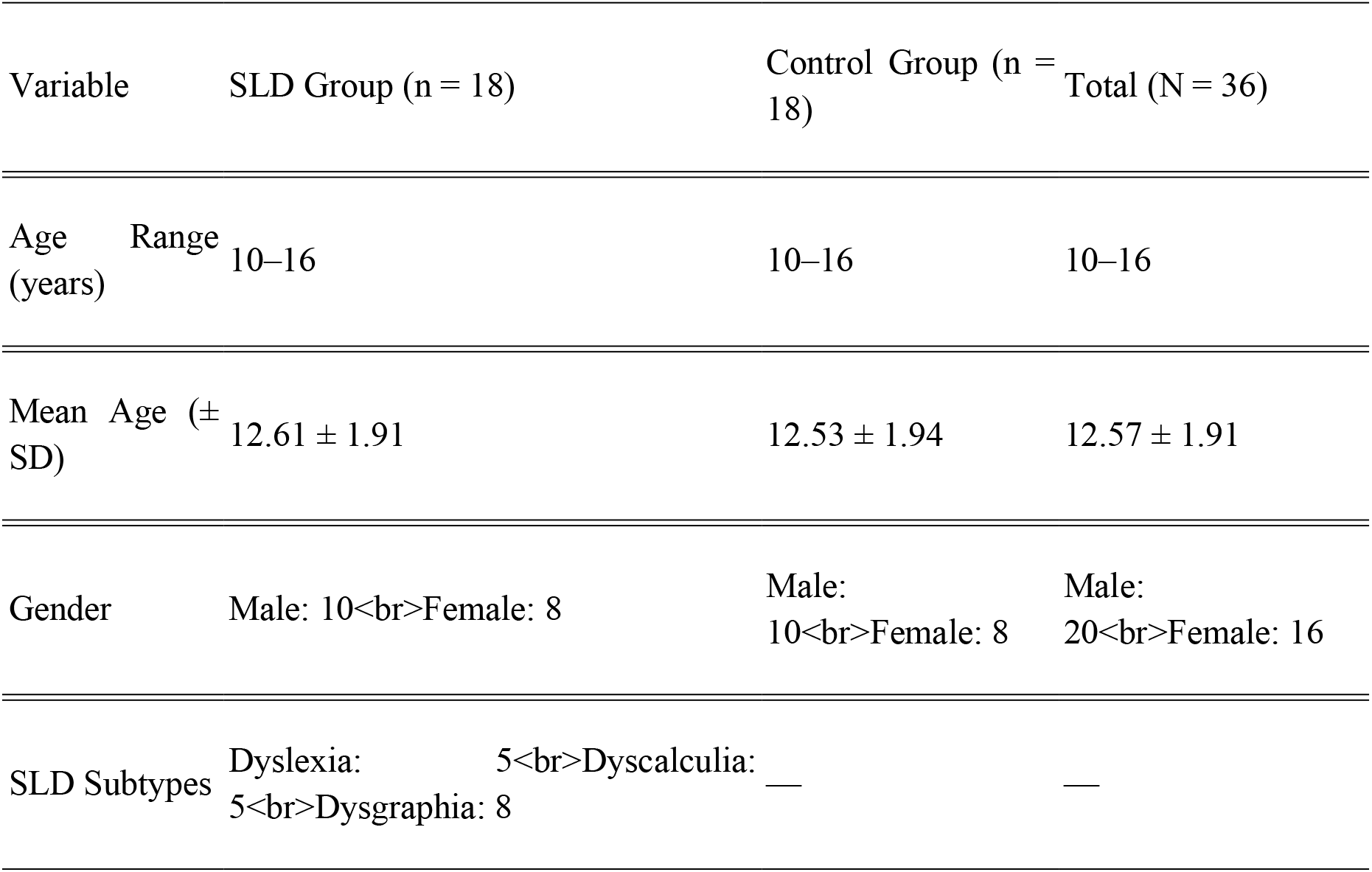
Demographic Characteristics of Study Participants.

### Accuracy

As expected accuracy increased reliably with larger intensity differences (β = 1.05, SE = 0.07, p < 1 × 10^−54^). Neither group (β = –0.27, SE = 0.27, p = 0.32) nor delay duration (β = –0.04, SE = 0.04, p = 0.32) exerted significant main effects. However, the interaction between group and stimulus intensity was significant (β = –0.75, SE = 0.08, p < 1 × 10^−19^), indicating reduced sensitivity to intensity differences in the SLD group compared with controls.

To further dissociate SLD subtypes, a second GLME included group type (dyslexia, dyscalculia, dysgraphia, control) as predictor. Across all subtypes, accuracy improved with increasing intensity differences (β = 1.04, SE = 0.06, p < 1 × 10^−62^). Yet interactions between subtype and intensity difference revealed attenuated slopes relative to controls: dyscalculia (β = –0.84, p < 1 × 10^−19^), dyslexia (β = –0.79, p < 1 × 10^−16^), and dysgraphia (β = –0.62, p < 1 × 10^−12^). No main effects of subtype or delay duration were observed (ps >0.05)

### Threshold & Sensitivity

Threshold values were higher in the SLD group compared with controls (β = 1.58, SE = 0.52, p = 0.0043), indicating that individuals with SLD required larger intensity differences to detect auditory changes. Examining SLD subtypes, only dyslexia showed a significant increase in threshold relative to controls (β = 2.51, p = 0.0022), while dyscalculia and dysgraphia exhibited non-significant or borderline effects.

Sensitivity values were significantly higher in the SLD group (β = 3.23, SE = 0.57, p < 1 × 10^−5^), suggesting enhanced psychometric discrimination. Subtype analyses revealed elevated sensitivity in dyscalculia (β = 4.99, p < 1 × 10^−7^) and dyslexia (β = 3.98, p < 1 × 10^−6^), with dysgraphia showing a smaller but significant increase (β = 1.66, p = 0.007). These results indicate distinct alterations in auditory discrimination thresholds and sensitivity across SLD subtypes.

### Reaction time

Linear mixed-effects models examined trial-level log-transformed reaction times (logRT) with fixed effects of accuracy, group, delay duration, and stimulus intensity difference. Participants in the SLD group were significantly slower than controls (β = 0.31, SE = 0.072, p < 0.001). Delay duration and interactions with the group did not significantly affect RTs (ps > 0.15). Reaction times increased slightly with larger stimulus intensity differences (β = 0.0044, SE = 0.002, p = 0.03), whereas accuracy had no significant effect on RTs (β = –0.015, p = 0.31).

A model including SLD subtypes indicated that all subgroups were slower than controls: dyscalculia (β = 0.30, p = 0.0046), dyslexia (β = 0.22, p = 0.034), and dysgraphia (β = 0.37, p < 0.00003). No interactions with delay or stimulus intensity were significant (ps > 0.18). These results suggest a general slowing in auditory working memory processing across SLD subtypes.

### Confidence level

Participants’ confidence ratings were high overall (M ≈ 74%). Confidence increased with larger stimulus intensity differences (β = 2.78, SE = 0.43, *p* < 1 × 10^−10^). No main effect of group was observed (β = –1.91, p = 0.75), but the interaction between group and intensity difference was significant (β = –1.61, SE = 0.61, *p* = 0.008), indicating reduced sensitivity of confidence to stimulus differences in the SLD group. Delay duration did not significantly affect confidence ratings (*p* = 0.27).

Examining SLD subtypes revealed that the interaction of type × stimulus intensity was significant only for dyslexia (β = –2.12, p = 0.02), with other subtypes showing no significant modulation. Including accuracy as a factor confirmed the negative group × intensity interaction (β = –1.27, p < 1 × 10^−6^) and showed a small negative effect of delay on confidence for correct responses (β = –0.88, p = 0.045). Overall, results indicate that while confidence generally tracks stimulus intensity differences, this sensitivity is attenuated in SLD, particularly in dyslexia

### Drift Diffusion Model (DDM)

To further disentangle latent cognitive mechanisms underlying performance differences, a hierarchical Bayesian diffusion decision model was fitted. Drift rate (μ), reflecting the efficiency of evidence accumulation, was significantly lower in the SLD group compared with controls (β = –0.07, 95% CI [–0.12, –0.01]), indicating slower accumulation of decision-relevant information. Boundary separation (a), indexing response caution, was significantly higher in the SLD group (β = 0.15, 95% CI [0.13, 0.18]), suggesting a more conservative decision strategy. Non-decision time (β = 0.03, 95% CI [0.02, 0.03]) did not differ between groups, indicating comparable sensory encoding and motor execution times. Starting point bias showed no reliable group effect (β = –0.01, 95% CI [–0.10, 0.08]). Together, these findings reveal that reduced drift rate and elevated boundary separation jointly contribute to altered decision dynamics in SLD participants.

## Discussion

The present study examined auditory working memory (AWM) performance and metacognitive monitoring in adolescents with Specific Learning Disorders (SLD). Across behavioral and computational measures, we observed clear alterations in perceptual sensitivity, decision dynamics, and confidence calibration relative to typically developing peers. These findings contribute to a growing body of evidence that SLD is not limited to domain-specific academic difficulties but also involves broader deficits in auditory discrimination and executive-level integration of sensory information.

At the level of task accuracy, SLD participants did not differ in overall baseline performance but demonstrated reduced sensitivity to incremental stimulus intensity. This suggests that while individuals with SLD can perform adequately under conditions of clear perceptual contrast, they struggle when discrimination demands are subtle. Notably, the impairment was observed across all subtypes, with dyscalculia and dyslexia showing the strongest effects. These results support prior neuroimaging findings of atypical temporoparietal activation during auditory discrimination in dyslexia (Faedda et al., 2019) and extend behavioral reports linking elevated auditory thresholds to phonological deficits (Snowling, 2020; McWeeny et al., 2024).

Analyses of discrimination thresholds and psychometric sensitivity highlighted a paradoxical pattern. SLD participants required larger intensity differences to reliably detect auditory changes, yet exhibited elevated sensitivity values compared with controls. This dissociation implies that perceptual encoding of auditory differences is intact—and in some cases exaggerated—but difficulties arise at the stage of translating this information into consistent decisions. This interpretation aligns with evidence that individuals with reading difficulties often display instability in auditory processing thresholds (Breadmore et al., 2024) and may experience inefficient executive filtering of sensory input (Czajeczny et al., 2023). Thus, sensory encoding may be preserved or even heightened, whereas the integration of perceptual cues into effective decisions remains inefficient.

Reaction time analyses further reinforced the presence of domain-general inefficiencies. Across subtypes, SLD participants responded more slowly than controls, independent of stimulus intensity or delay duration. This generalized slowing is consistent with electrophysiological findings of delayed P300 latencies in SLD (Kashani & Tafti, 2018) and with behavioral evidence of prolonged processing times linked to executive dysfunction (Shyam & Venkatesan, 2021). Together, these data suggest that deficits in auditory discrimination tasks are not confined to sensory processing but reflect broader limitations in cognitive efficiency and decision-making speed.

Metacognitive analyses revealed an additional layer of impairment. Although baseline confidence was comparable between groups, individuals with SLD exhibited reduced sensitivity of confidence ratings to stimulus differences. In other words, their subjective certainty did not scale appropriately with objective task difficulty, particularly among participants with dyslexia. This dissociation between objective performance and subjective monitoring converges with recent work showing that students with learning disabilities benefit less from confidence-based self-monitoring interventions (Okur & Aksoy, 2025; Daniel et al., 2022). These findings suggest that impaired calibration between confidence and performance may undermine effective self-regulation in academic settings.

Drift diffusion modeling provided mechanistic insight into these behavioral patterns. The SLD group showed lower drift rates, indicating slower evidence accumulation, and larger boundary separation, reflecting more conservative response strategies. These findings explain the combined pattern of reduced perceptual sensitivity, slower responses, and altered confidence calibration: while sensory encoding is available, decision-level integration proceeds more slowly and cautiously, creating a bottleneck between perception and action. Importantly, non-decision times and starting point biases did not differ between groups, confirming that the core deficits lie in evidence integration rather than sensory encoding or motor execution. This is consistent with earlier modeling work in dyslexia showing reduced drift rates during perceptual tasks (Banai & Ahissar, 2006) and theoretical accounts linking conservative strategies to uncertainty in phonological representations (Snowling, 2020).

Several limitations should be acknowledged. The relatively small sample size and cross-sectional design restrict the generalizability of our findings and preclude causal inferences. Moreover, the heterogeneity within SLD subtypes may have introduced variability that was not fully captured in the present analyses. In addition, our study relied exclusively on behavioral and computational measures without integrating neurophysiological data such as EEG or fMRI, which could provide direct evidence of underlying neural mechanisms. Future research should therefore employ larger, longitudinal samples and multimodal approaches to clarify developmental trajectories and brain– behavior relationships in auditory working memory. From an applied perspective, intervention studies are needed to evaluate whether training programs that target metacognitive monitoring and decision-making strategies can improve academic outcomes in adolescents with SLD.

In summary, this study demonstrates that adolescents with SLD exhibit a distinctive profile of auditory working memory dysfunction. Their difficulties are not solely perceptual but extend to metacognitive and decisional domains, marked by reduced discrimination sensitivity, generalized cognitive slowing, and attenuated confidence calibration. By combining behavioral, psychometric, and computational approaches, we identify a decision-level bottleneck that may explain why individuals with SLD struggle to translate sensory input into efficient and confident judgments. These insights underscore the importance of targeting not only phonological training but also metacognitive monitoring and decision-making strategies in educational and clinical interventions for SLD.

